# Maternal choline supplementation lessens the behavioral dysfunction produced by developmental manganese exposure in a rodent model of ADHD

**DOI:** 10.1101/2023.06.23.546356

**Authors:** Shanna L. Howard, Stephane A. Beaudin, Barbara J. Strupp, Donald R. Smith

## Abstract

Studies in children have reported associations between elevated manganese (Mn) exposure and ADHD-related symptoms of inattention, impulsivity/hyperactivity, and psychomotor impairment. Maternal choline supplementation (MCS) during pregnancy/lactation may hold promise as a protective strategy because it has been shown to lessen cognitive dysfunction caused by numerous early insults. Our objectives were to determine whether (1) developmental Mn exposure alters behavioral reactivity/emotion regulation, in addition to impairing learning, attention, impulse control, and sensorimotor function, and (2) MCS protects against these Mn-induced impairments. Pregnant Long-Evans rats were given standard diet, or a diet supplemented with additional choline throughout gestation and lactation (G3 - PND 21). Male offspring were exposed orally to 0 or 50 mg Mn/kg/day over PND 1–21. In adulthood, animals were tested in a series of learning, attention, impulse control, and sensorimotor tasks. Mn exposure caused lasting dysfunction in attention, reactivity to errors and reward omission, learning, and sensorimotor function, recapitulating the constellation of symptoms seen in ADHD children. MCS lessened Mn-induced attentional dysfunction and partially normalized reactivity to committing an error or not receiving an expected reward but provided no protection against Mn-induced learning or sensorimotor dysfunction. In the absence of Mn exposure, MCS produces lasting offspring benefits in learning, attention, and reactivity to errors. To conclude, developmental Mn exposure produces a constellation of deficits consistent with ADHD symptomology, and MCS offered some protection against the adverse Mn effects, adding to the evidence that maternal choline supplementation is neuroprotective for offspring and improves offspring cognitive functioning.

**Highlights:**

1. Developmental Mn exposure causes lasting dysfunction consistent with ADHD symptomology.
2. Maternal choline supplementation (MCS) protects against Mn-induced deficits in attention and behavioral reactivity.
3. MCS in control animals produces lasting benefits to offspring in learning, attention, and error reactivity.
4. These data support efforts to increase choline intake during pregnancy, particularly for individuals at risk of neurotoxicant exposure.

## 1. Introduction

Attention Deficit Hyperactivity Disorder (ADHD) is the most prevalent neurodevelopmental disorder among children, affecting ∼5-7% of youths up through age 18 years, and ∼2–3% of adults^1^. ADHD is characterized by symptoms of inattention, impulsivity/hyperactivity, emotion dysregulation, and in some cases psychomotor deficits^2–4^. Notably, emotion dysregulation has been shown to have a greater impact than inattention and hyperactivity on overall well-being and self-esteem, including peer relationships, academic performance and occupational attainment^5,6^. While the cause(s) of ADHD and related symptoms remains poorly understood, there is an emerging body of epidemiological evidence showing that developmental environmental insults, such as exposure to elevated manganese (Mn) levels, is associated with increased risk of ADHD and related symptoms in children and adolescents^7,8,17,9–16^. Although these epidemiological studies have played a key role in raising concerns about elevated Mn exposure, they are unable to establish causality, in part because of challenges in accurately measuring Mn exposure, and quantifying and controlling for potential confounding factors.

Our prior rodent model studies have shown that developmental Mn exposure can cause lasting ADHD-like impairments in attention, impulse control, and sensorimotor function. In addition, these deficits are associated with hypofunctioning of the catecholaminergic system in the prefrontal cortex and striatum - brain regions that in part mediate attention, impulse control, emotion regulation, and sensorimotor function^18–25^. While our rat model of developmental Mn exposure recapitulates many of the features of ADHD, it remains unclear whether Mn exposure also causes lasting disruption of behavioral reactivity/emotion regulation. This is a significant knowledge gap, given the central role that emotion dysregulation plays in ADHD symptomology and quality of life for children with ADHD^3–5^, and the fact that Mn exposure is a risk factor for ADHD diagnosis^7^.

While therapies such as methylphenidate (Ritalin) have proven to be effective for children diagnosed with ADHD, there are no established treatments available for children exposed to high levels of Mn during development. A number of nutritional interventions, including maternal supplementation with choline, folic acid, genistein, and L-methionine have been shown to mitigate the effects of a variety of environmental insults^26,27^. In particular, maternal choline supplementation (MCS) has been shown to lessen cognitive dysfunction caused by prenatal alcohol exposure^28,29^, maternal stress^30^, and Down syndrome^31–33^ in animal models, to normalize some gene expression and epigenetic molecular alterations caused by perinatal alcohol exposure^34,35^, and to mitigate some of the detrimental effects of prenatal alcohol exposure in children of alcohol-using mothers^36,37^. In addition, MCS has been shown to improve cognitive function of typically developing children^38–40^ and animal models^41–49^. These benefits of MCS may reflect, in part, the growing evidence that the amount of choline most commonly consumed by pregnant women – and pregnant laboratory animals – is insufficient to meet the increased demand for choline during fetal and neonatal development^50–52^. Physiological demands for choline increase markedly during pregnancy due to choline’s numerous roles in fetal development, including as a precursor biomolecule for acetylcholine, phosphatidylcholine, sphingomyelin, and betaine^38,53,54^. However, over 90% of pregnant women consume less than the Adequate Intake (AI) for choline^40,50,51,55^ – an amount which itself appears to be insufficient to meet the demands of pregnancy^38–40^.

The present study used our rodent model of early childhood environmental Mn exposure to test two hypotheses: (1) That developmental Mn exposure causes lasting alterations in behavioral reactivity/emotion regulation (assessed via the reaction to committing an error or not receiving an expected reward) in addition to the deficits in attention, impulse control, learning, and sensorimotor function – all areas of dysfunction seen in children with ADHD; and (2) that supplementing the maternal diet with additional choline during pregnancy and lactation will offer protection against the adverse effects of early postnatal Mn exposure on attention, impulse control, learning, sensorimotor function, and behavioral reactivity (if shown). Collectively, these findings will add to our understanding of the constellation of effects produced by developmental Mn exposure as a risk factor for ADHD, and they will provide evidence to evaluate the efficacy of MCS to protect against Mn-induced dysfunctions.

## 2. Materials and Methods

### 2.1 Subjects

Sixty-four (64) male Long-Evans rats were used for neurobehavioral assessment. Additional littermates were used for analyses of tissue Mn, choline, and choline metabolites. Subjects were born in-house from 16 nulliparous timed-pregnant dams (Charles River; gestational age 3; 8-9 weeks old). Twelve to 24 hours after parturition (designated PND 1, birth = PND 0), litters were weighed, sexed, and culled to eight pups per litter such that each litter was composed of as many males as possible (2-8) and the remainder females. Each treatment group contained n = 16 animals, and no more than 1-2 males per litter were assigned to a particular treatment group. At PND 22, all pups were weaned and pair-housed with an animal of the same treatment group and maintained on a reversed 12:12 hour light/dark cycle. Animals were housed in polycarbonate cages at a constant temperature of 21 ± 2°C. All aspects of behavioral testing and feeding were carried out during the active (dark) phase of the animals’ diurnal cycle. All animal care and treatments were approved by the institutional IACUC (protocol Smitd1803) and adhered to National Institutes of Health guidelines set forth in the Guide for the Care and Use of Laboratory Animals^56^.

### 2.2 Maternal choline supplementation

Pregnant dams were randomly selected to receive one of two concentrations of choline chloride in their diet (Dyets AIN-76A Purified Rodent Diet), both of which were provided *ad libitum* during gestation and lactation, from G3 until PND 21. The control purified diet (unsupplemented) contained 1.1 g choline/kg, whereas the supplemented purified diet contained 5.0 g choline/kg; both are reported by the manufacturer to contain 54 mg Mn/kg. Upon weaning on PND 21, all pups were provided Irradiated Teklad Global Soy Protein-Free Extruded Rodent Diet 2920, reported by the manufacturer to contain 80 mg Mn/kg and 1.2 g choline/kg. See Supplemental Figure S1 for a timeline of maternal choline supplementation, Mn dosing, and behavioral testing.

### 2.3 Manganese exposure

Neonate animals were orally exposed to 0 or 50 mg Mn/kg/day in a vehicle solution (5% w/v stevia in Milli-Q water) from PND 1–21, as in our prior studies^18–24^. Briefly, Mn was delivered once per day directly into the mouth of each pup (∼15-25 μL/dose) using a micropipette fitted with a flexible polyethylene pipette tip. This Mn exposure regimen is relevant to children exposed to elevated Mn via drinking water, diet, or both, as pre-weaning exposure to 50 mg Mn/kg/day produces relative increases in Mn intake that approximates the relative increases reported in infants and young children exposed to Mn-contaminated water or soy-based formulas (or both)^20,22^.

### 2.4 Testing apparatus

Sixteen identical automated 5-Choice Serial Reaction Time Task (5-CSRTT) testing chambers fitted with odor delivery systems (#MED-NP5L-OLF, Med Associates, Inc., St. Albans, VT) were used to assess specific cognitive processes, including learning, focused and selective attention, inhibitory control, and behavioral reactivity, as described previously^20,21,57^. Briefly, each testing chamber contained a curved wall equipped with five 2.5 × 2.5 cm response ports fitted with light-emitting diodes that served as the visual cue, an infrared beam to register nose pokes, and pneumatic inlet and vacuum outlet ports to introduce and remove air-based odor distractors. Opposite the response wall was the food magazine wall that contained a 45 mg food pellet reward dispensing port fitted with an infrared beam to register nose pokes; each unit also contained a small house light. The entire testing chamber was enclosed in a sound attenuating cubicle. Each animal was assigned to one of the 16 testing chambers, and each chamber was balanced for the four treatment conditions.

### 2.5 Behavioral testing

Animals were mildly food restricted starting on PND 45 in preparation for behavioral testing, with water available *ad libitum* throughout behavioral assessment. Animals were provided food for 2 hours each day immediately after behavioral testing, so that their body weights were maintained at ∼90–95% of free-feeding weights. All rats were weighed and tested 6 days/week throughout training and testing.

Behavioral assessment occurred during the active (dark) period of the diurnal cycle at the same time each day and in the same chamber for each individual rat. A daily test session consisted of 150 trials or 50 min, whichever came first. Each trial sequence was initiated by a nose-poke in the food magazine port, which was followed by a 3 sec turnaround time to allow the animal to reorient from the food magazine wall to the response wall; trial onset began after the 3 sec turnaround time. All behavioral testing was conducted by individuals blind to the treatment condition of the subjects.

Behavioral testing began on ∼ PND 50. Briefly, testing began with food magazine and nose-poke training for 11 days, followed by a 5-choice visual discrimination learning task in which the visual cue was presented immediately after trial onset (i.e., no pre-cue delay), and stayed illuminated for 15 sec (for more detail, see Beaudin et al.^20^). Animals stayed on this task until they reached the criterion of 80% correct trials on 2 of 3 consecutive testing days. Following this task, animals were administered a second visual discrimination task with a fixed 1 sec cue duration and no pre-cue delay for 5 days. This training was followed by a series of attention and behavioral reactivity tasks as described below (see Supplemental Figure S1 for the timeline of testing).

5-CSRTT assessment is conducted via a series of progressively more difficult tasks, described in greater detail below, that challenge the ability of animals to sustain attention, inhibit premature responses, and ignore olfactory distractors while maintaining focus on the predictive visual cue. Recorded response types for all 5-CSRTT tests included the following: 1) premature responses (responses made after trial onset but before presentation of the visual cue; assessed in the first focused attention, second focused attention, and selective attention tasks); 2) correct responses (responses made to the correct port following presentation of the visual cue); 3) incorrect responses (responses made to the incorrect port following presentation of the visual cue); and 4) omissions (failure to respond within 15 sec after visual cue presentation). Premature and incorrect responses and omission errors were not rewarded and were immediately followed by a 5 sec time-out, in which the house light was turned off for 5 sec. In addition, the latency (sec) for correct and incorrect responses was recorded (response latency), as was the latency (sec) to retrieve the food pellet reward following a correct response (collection latency). Perseverative responses, defined as additional nose pokes into the response port after a correct response, were also recorded. The calculated response outcomes were: (1) %Correct, calculated as number of correct responses/(correct + incorrect + premature + omissions) × 100; (2) %Incorrect, calculated as above but with incorrect responses in the numerator; (3) %Premature, calculated as above but with premature responses in the numerator; and (4) %Omissions, calculated as above but with omissions in the numerator. Additionally, we assessed attentional accuracy via a %Accurate outcome, calculated as number of correct responses/(correct + incorrect) × 100. The number of perseverative responses per correct response was calculated simply as #perseverative responses/#correct responses.

### 2.6 Focused attention tasks

Focused attention can be defined as the ability to maintain attentional focus on a specific task or stimulus (e.g., a visual cue). Two successive focused attention tasks (FA1 and FA2) were administered over ∼PND 85-103, and PND 104-121, respectively, following completion of the visual learning discrimination task. Both focused attention tasks assessed the ability of animals to detect and respond to a visual cue presented unpredictably in time and location (one of the five response ports). The first focused attention task used variable pre-cue delays of 0, 1, 2, or 3 sec and a fixed visual cue duration of 0.7 sec and was administered for 15 daily sessions. The second focused attention task included variable pre-cue delays of 0, 3, 4, or 5 sec and variable visual cue durations of 0.5 or 0.7 sec and was administered for 15 sessions.

### 2.7 Selective attention task with olfactory distractors

Selective attention can be defined as the ability to maintain a behavioral or cognitive set in the face of distracting or competing stimuli. The selective attention task (SAT) with olfactory distractors was administered across 12 sessions, from PND 129-140. In this task, the pre-cue delay varied between 3 and 4 sec, with the two delays balanced across the trials within each test daily session, and the cue duration was 0.5 sec. On one third of the trials in each session, an olfactory distractor was presented 1 or 2 sec after trial onset (i.e., 1–3 sec before the visual cue, depending on the pre-cue delay). The delay between odor distractor presentation and visual cue presentation (1-3 sec) is referred to as the odor-to-cue interval (OtoC) and was included in statistical analysis. The nine different olfactory distractors were made from liquid odorants (McCormick & Company, Inc.) diluted in propylene glycol, and delivered into an incorrect response port as scented air. The SAT was preceded by a baseline task (three sessions), administered from PND 126-128. This task was identical to the SAT, except it did not include the presentation of olfactory distractors.

### 2.8 Assessment of behavioral reactivity and compulsiveness in the reward omission task

Behavioral reactivity/emotion regulation was assessed by measuring the animal’s performance on trials following an error or not receiving an expected reward, areas where children with ADHD have been shown to differ from typically developing children^3,4^. Reactivity to committing an error was assessed in two tasks: (1) The second focused attention task, and (2) a reward omission task. In this latter task, a 0.7 sec visual cue was presented immediately at trial onset, and the food pellet reward was randomly omitted on three out of 10 correct trials. It is notable that the consequences of committing an error versus a reward omission are very different for the animal. Upon committing an error (in all tasks), the rat receives a 5 sec ‘time-out’ during which the house light turns off and the animal sits in darkness, and no pellet reward is provided. In contrast, on a reward omission trial the house light remains on and all other trial conditions/cues remain unchanged from a normal rewarded correct trail. Attentional performance on the reward omission task was assessed as a function of whether the preceding trial was either (1) correct (and rewarded), (2) incorrect (including omission errors), or (3) correct but food reward omitted. We also assessed perseverative responses in the reward omission task as a measure of compulsivity, consistent with prior studies^58,59^. Perseverative responses are additional nose-pokes into the correct port following a correct response. Animals were tested in the reward omission task for 12 sessions from PND 175-187.

### 2.9 Montoya staircase task

The Montoya staircase test was used to assess sensorimotor function, as described previously by us^19^. Performance on the staircase test is a sensitive measure of skilled forelimb function directly relevant to pediatric Mn studies. This test taps essential sensorimotor functions required to successfully reach and grasp small food pellets from the descending steps of the staircase, including advancing the limb over the food, opening the digits in preparation for grasping, grasping and manipulating the food, and withdrawing the paw to place the food in the mouth^60^. Optimal performance on this test requires several sensorimotor skills that are similarly assessed in human studies, including forelimb movements and digit flexion for manipulating objects. Thus, the staircase test measures aspects of fine psychomotor skills that are known to be disrupted in children with a history of developmental Mn exposure^8,61^.

Eight identical Plexiglas staircase devices were used, modeled after the original design of Montoya et al.^62^. Each device contained a platform and a left-and-right side staircase. The staircases contained 6 descending steps on each side, and each step was baited with three 45 mg food pellets for a total of 36 pellets available at the start of a session. To determine the step of origin and final location of each pellet, pellets were color coded with a unique color per step level^63^.

The staircase study was conducted over 1 month, beginning on PND 85. Each staircase session occurred immediately following operant testing every day. Staircase training occurred for the first 11 days. During training, rats retrieved the food pellets from the descending steps of the staircases until all subjects reached asymptotic performance of total pellets retrieved. The next 19 days comprised the testing phase of the task. Throughout training and testing, each subject was given one 10-min trial per day, 6 days per week. After each trial, the rat was removed, and the final distribution of colored pellets was quantified per step and side. The order and time of testing were balanced by treatment and remained the same each day for each rat.

Skilled forelimb performance was evaluated step-by-step for (1) the number of pellets taken (i.e., pellets that were grasped and removed from their step of origin), (2) the number of pellets eaten (pellets that were taken and consumed), (3) the corresponding percent of success (the ratio of the number of pellets eaten/pellets taken x 100), (4) the number of pellets misplaced (all the pellets that were grasped but ended up on a different step level than their step of origin), and (5) the number of pellets lost (pellets that ended up on the floor of the apparatus, out of reach of the animal).

### 2.10 Tissue Mn and choline/choline metabolite levels

For blood and brain Mn levels, tissue samples were collected from weaned pups on PND 24, and from the behaviorally tested animals on PND 202. Briefly, animals were euthanized via CO_2_ asphyxiation, and whole blood was immediately collected from the left ventricle of the surgically-exposed heart and stored in EDTA Vacutainers at − 20 °C for analyses. Whole brain was immediately removed, and the hind-brain region collected and stored at −80 °C for Mn concentration determinations. Blood and brain Mn concentrations were determined by inductively coupled plasma–mass spectrometry (Thermo Element XR), as previously described^18,20^. The analytical detection limit for Mn was 0.01 ng/mL.

Choline/choline metabolite levels were measured in plasma, brain, and/or liver tissues from dams at weaning (plasma and liver), and/or from the same PND 24 animals (liver and hindbrain) used for Mn analyses described above. Briefly, concentrations of free choline, betaine, dimethylglycine, methionine, and trimethylamine N-oxide were measured by liquid chromatography tandem mass spectrometry according to the method of Holm et al.^64^, while phosphocholine, glycerophosphocholine, phosphatidylcholine, acetylcholine, and lysophosphatidylcholine were measured by liquid chromatography/mass spectroscopy according to the method of Koc et al.^65^ with modifications^66^.

### 2.11 Statistical methods

The behavioral data were modeled using structured covariance mixed models. Fixed effects included in the models were the between-subject factors MCS (two levels corresponding to the unsupplemented and MCS groups), and Mn exposure treatment (two levels corresponding to the 0 and 50 mg/kg/d groups), and the within-subject factors pre-cue delay, cue duration, session block, distraction condition, and/or previous trial outcome, depending on the outcome analyzed. In all models, rat was included as a random effect to account for correlations within observations from the same animal. Statistical tests used the Kenward-Roger correction to improve estimation of the variance-covariance of the fixed effects parameters. Plots of residuals by experimental condition were used to examine the assumption of homogeneity. Random effects with high variance in the residuals across the levels of the within-subject factor (e.g., distraction condition) were added to the model to achieve homogeneity if necessary. The distribution of each random effect was inspected for approximate normality and presence of influential outliers. The significance level was set at p ≤ 0.05, and p-values between 0.05 and 0.10 were considered to be trending towards significance; trending effects are presented if they aid in clarifying the nature of the Mn and/or MCS effects. Significant main effects or interaction effects were followed by single-degree of freedom contrasts to better understand the nature of the interactions, using the Student’s t-test for pairwise comparisons of least squared means. Analyses were conducted using SAS (version 9.4) for Windows. For tissue Mn and choline/metabolite data, analyses were performed using one-way (for dams) or two-way (for offspring) analysis of variance and Tukey’s post hoc test for pairwise comparisons, using JMP (version 16.0; SAS Institute, Inc.).

## 3. Results

For the 5-CSRTT measures, we focus primarily on outcomes of %Premature, %Correct, and %Accurate responses because our prior studies have shown that these measures best differentiated control vs Mn animals^20,21^ and did so again in this study. Note that in general, we graphically depict the data for all four treatment groups (defined by diet and Mn exposure), regardless of whether the interaction of Mn and MCS was significant, due to our *a priori* hypotheses relating to the magnitude of the Mn effect under each of the two diet conditions.

### 3.1 Visual learning

#### Visual learning in the 5-choice task is not altered by Mn or MCS

Visual learning (i.e., learning to make a nosepoke into the illuminated port) is foundational to the subsequent, more challenging 5-CSRTT tasks. Here, there was no effect of early postnatal Mn exposure or MCS on visual learning. Specifically, there was no main effect of Mn or MCS on trials to criterion or days to criterion, and no Mn x MCS interaction for these two outcomes (p’s >0.295). Moreover, there also was no effect of Mn or MCS on the percentage of animals per treatment group failing to reach the learning criterion. The lack of treatment group differences in learning the task criterion indicates that visual acuity and motivation did not differ between groups and therefore cannot account for group differences seen in subsequent 5-CSRTT tasks.

### 3.2 Impulsivity

#### Developmental Mn exposure did not affect impulse control, but did impair the animals in learning to wait for the visual cue, and this effect is not rescued by MCS

We evaluated %Premature responses as a measure of inhibitory control in the two focused attention tasks and the selective attention task.

##### In the first focused attention task, Mn exposure impairs learning to withhold premature responses, and MCS does not alter this effect

In the first focused attention task, there is a significant Mn x session block interaction [F(2, 435.8) = 11.07, p<0.0001] for %Premature responses. Whereas both control and Mn groups commit a relatively high percentage of premature responses at the start of the task (∼ 50%, since this was the first task in which animals faced longer pre-cue delays), the Mn animals improve less rapidly than controls. As a result, the Mn animals commit significantly more premature responses than controls in session block 3 (p=0.0084; Figure 1a1). There is also a significant MCS x session block interaction [F(2, 435.8) = 17.06, p<0.0001], but none of the contrasts achieved significance (this two-way interaction is not plotted, but all four groups are presented in figure 1A.) The three-way interaction of MCS, Mn and block was not significant; p = 0.69).

**Figure 1.**
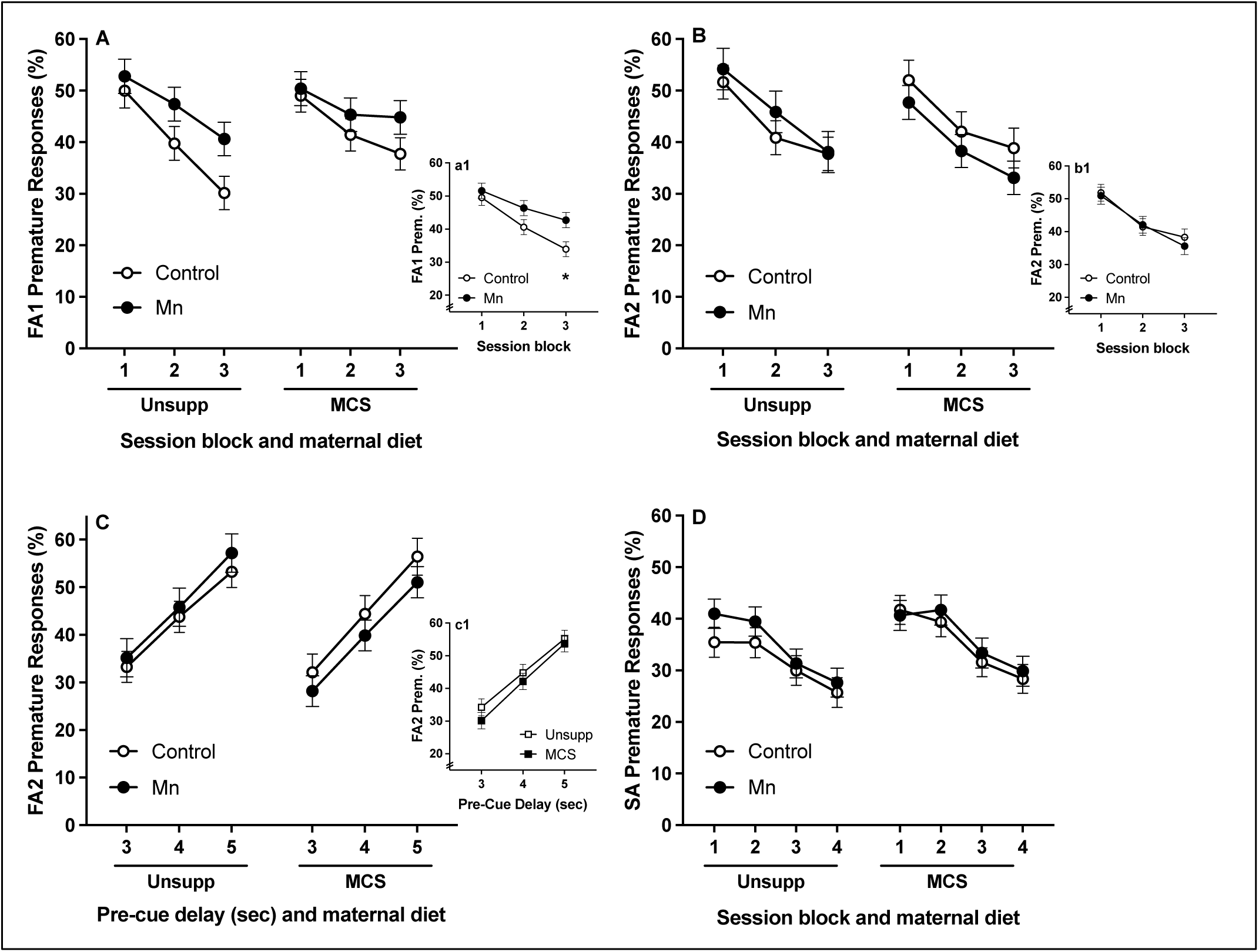
Effects of Mn and MCS on premature responses (impulsivity) in the focused attention 1 task (panel A), the focused attention 2 task (panels B and C), and the selective attention task (panel D), as a function of session block or pre-cue delay (sec). Sub-panels a1, b1, and c1 depict significant two-way interactions of Mn x session block or MCS x pre-cue delay, respectively. Mn exposure impaired the rate of learning to wait for the cue. MCS did not alter premature responses generally, although there is a suggestion that MCS lessened group differences in the selective attention task. * indicates p ≤ 0.05 versus controls. Data are lsmeans ± SEM (n=14-16/group).

##### In the second focused attention task, neither Mn nor MCS significantly alter premature responses

In the subsequent second focused attention task, neither main effect of Mn (p = 0.75) nor MCS (p = 0.39) was significant. However, there is a Mn x session block interaction [F(2, 933.7) = 5.53, p = 0.0041], although there are no significant contrasts between control and Mn animals in any session block (p’s >0.40; Figure 1b1) (see also Figure 1B, which depicts differences between Mn and controls under each of the two diet conditions; the Mn x MCS x block interaction p = 0.40). There is also a MCS x pre-cue delay interaction [F(2, 932.8) = 3.18, p = 0.0422], though again, none of the specific contrasts between unsupplemented and MCS animals reached significance at any pre-cue delay (p’s > 0.21, Figure 1c1). Differences between Mn and controls under each of the two diet conditions is presented in Figure 1C (the three-way interaction of MCS, Mn and pre-cue delay was not significant; p = 0.24).

##### In the selective attention task, MCS lessens the difference in premature responses between treatment groups in early session blocks

In the selective attention task with olfactory distractors, there is a Mn x MCS x session block interaction for %Premature responses [F(3, 1665) = 4.44, p = 0.0041], although none of the specific contrasts between treatment groups or diet conditions reached significance for any session block (p’s > 0.12, Figure 1D). Nonetheless, the significant three-way interaction appears to reflect the fact that the difference in premature responses between Mn and control groups in the first two session blocks is greater for unsupplemented animals than for the MCS animals, suggesting that MCS lessened the impulsivity caused by developmental Mn exposure.

### 3.3 Attention

#### Mn impairs focused attention and MCS lessens this effect

As described earlier, the two focused attention tasks were used to assess whether Mn exposure alters the ability of animals to maintain attentional focus, and whether the Mn impairment was lessened by MCS.

##### In the first focused attention task, Mn exposure impairs attention, and MCS improves learning of the control group

In the first focused attention task, there is a significant Mn x session block interaction for %Correct responses [F(2, 595.6) = 4.12, p = 0.0167]. This reflects that performance in the Mn animals improves more slowly than control animals across session blocks, such that the Mn animals have significantly fewer %Correct responses than the controls in session blocks two and three (p=0.007 and 0.002, respectively, Figure 2a1) (see Figure 2A, which depicts differences between Mn and controls under each of the two diet conditions; the three-way interaction p = 0.22).

**Figure 2.**
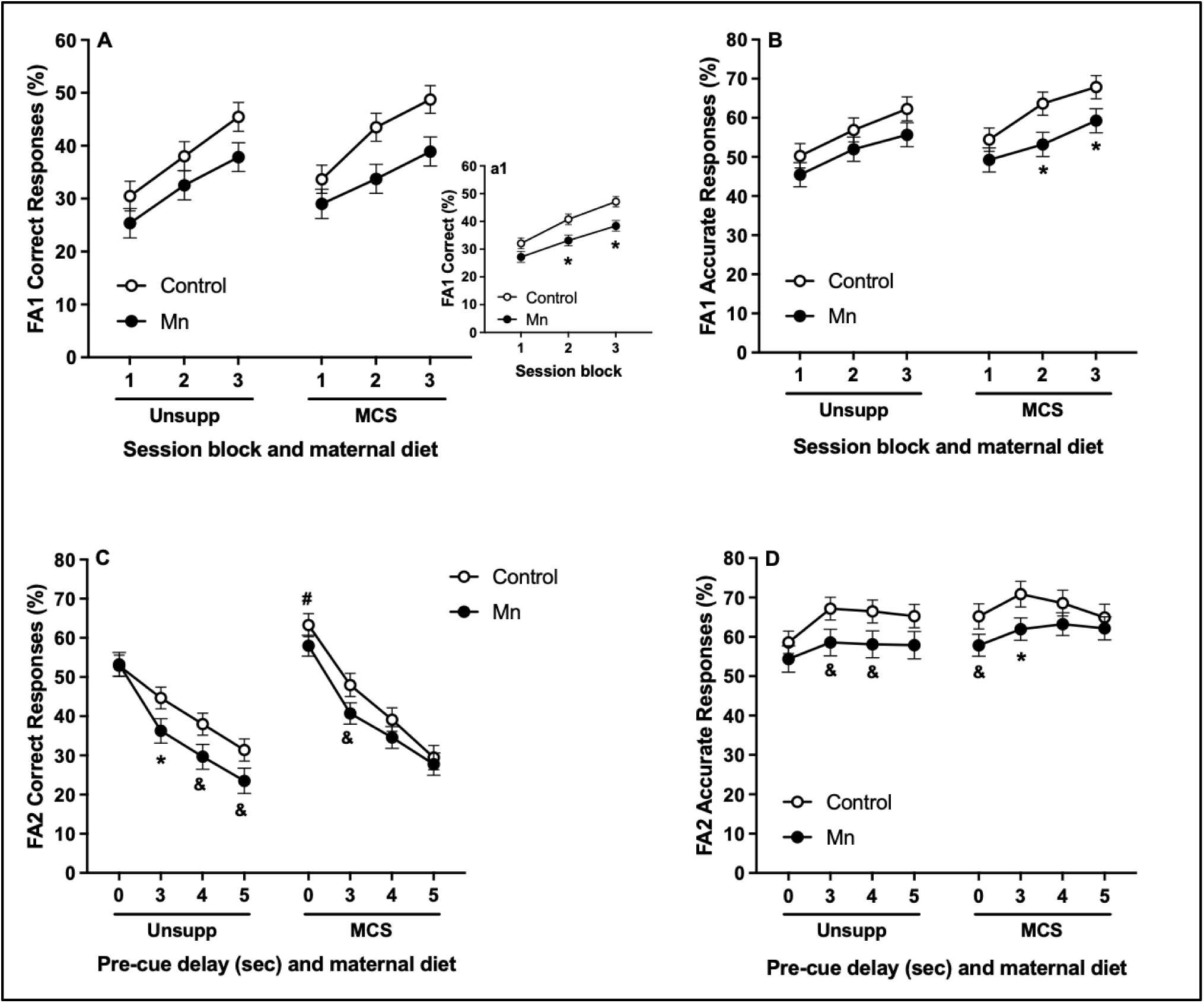
Effects of Mn and MCS on %Correct and %Accurate responding in the two focused attention tasks. Panels A and B depict performance in the focused attention 1 task as a function of session blocks; sub-panel a1 depicts the significant two-way interaction of Mn x session block. Panels C and D depict performance in the focused attention task 2, as a function of the pre-cue delay (sec). Mn animals exhibit an impairment in focused attention, and MCS affects attention in both groups, but in different ways. * and & indicate p ≤ 0.05 and p ≤ 0.10 versus controls, respectively; # indicates p ≤ 0.05 versus unsupplemented animals. Data are lsmeans ± SEM (n=14-16/group).

Consistent with the %Correct outcome noted above, there is a significant Mn x session block interaction for %Accurate responses [F(2, 601.3) = 3.33, p = 0.0365; not shown], and a trending three-way interaction of Mn x MCS x session block for %Accurate responses [F(2, 601.3) = 2.58, p = 0.0769] (Figure 2B). This three-way interaction reflects the fact that, while all groups improve in attentional accuracy across session blocks (p’s<0.0014), the difference in %Accuracy of Mn and control animals was greater in the MCS condition than the unsupplemented condition. This appears to be because the controls benefitted more from MCS than the Mn animals, in terms of improvement in %Accuracy across session blocks. As a result, under MCS, a Mn deficit in %Accuracy emerges under session blocks two and three (p’s = 0.018 and 0.049, respectively; Figure 2B).

##### In the second focused attention task, MCS improves performance of both the Mn and control animals, but in different functional domains

In the second focused attention task there is a significant Mn x MCS x pre-cue delay interaction for

%Correct responses [F(3, 1277) = 7.32, p < 0.0001] (Figure 2C). For animals on the unsupplemented diet, the Mn-exposed rats have significantly lower %Correct responses than controls at the 3 sec delay (p = 0.0477), and this Mn deficit is also trending at the 4 and 5 sec delays (p’s = 0.0518 and 0.0699, respectively). However, in the MCS animals, the Mn deficit at the 3 sec delay is only trending (p = 0.0762), and the deficit is no longer present under the 4 and 5 sec pre-cue delays (p’s = 0.27 and 0.69, respectively). Thus, the Mn x MCS x pre-cue delay interaction reflects the fact that the differences between control and Mn animals at the longer delays is smaller under MCS than under the unsupplemented diet. Additionally, control animals that received MCS have significantly more correct responses at the 0 sec delay compared to unsupplemented controls (p = 0.0113), reflecting a benefit of MCS to control animals for trials when the cue was presented immediately at trial onset, with no pre-cue delay.

We also assessed attentional accuracy (as %Accurate responses) for the second focused attention task, and again observed a significant Mn x MCS x pre-cue delay interaction [F(3, 1271) = 3.95, p = 0.0081] (Figure 2D), reflecting a pattern in attentional performance similar to what was seen with the %Correct outcome described above. Under the unsupplemented diet, there is a trending Mn deficit at the 3 and 4 sec pre-cue delays (p’s = 0.058 and 0.067, respectively). In contrast, under MCS, this Mn deficit is still present at the 3 sec delay (p = 0.043), but not at the 4 sec pre-cue delay (p = 0.23). While there was no significant Mn deficit at the 5 sec pre-cue delay under the unsupplemented diet, MCS also appears to reduce the difference between control and Mn animals at this longer delay (p = 0.11 under the unsupplemented diet and 0.53 under MCS; Figure 2D).

While there was a significant benefit of MCS to controls at the 0 sec pre-cue delay for %Correct responses (p = 0.0113), for %Accurate this benefit did not reach significance (p = 0.129). There was, however, a trending Mn deficit that emerged at the 0 sec delay under MCS (p = 0.09, Figure 2D).

Collectively, this evidence suggests that the lasting attentional deficits caused by developmental Mn exposure persist into the second focused attention task. Additionally, MCS improves performance of the controls but not the Mn animals for trials with a 0 sec pre-cue delay, whereas it reduces the difference between Mn and control animals at the longer (more challenging) delays. The MCS benefit derived by the controls at the 0 sec delay appears to be due to a lessened reaction to committing an error on the prior trial (discussed below; Figure 4), whereas the benefit derived by the Mn animals at the longer delays appears to reflect improved attention.

### MCS normalizes the performance of the Mn rats on distraction trials in the selective attention task

To determine the effects of Mn exposure and MCS on selective attention, we assessed animal performance on a selective attention task with olfactory distractors randomly presented either 1, 2, or 3 sec prior to the onset of the visual cue (referred to as the odor-to-cue interval, ‘OtoC’).

#### Mn causes attentional dysfunction and MCS reduces the difference between Mn and control animals on trials with olfactory distractors

For %Correct responses in the selective attention task, there was a significant Mn x OtoC interaction [F(3, 239.2) = 2.94, p = 0.0339], where the Mn animals were more impaired relative to controls for trials without a distractor (p = 0.0096; Figure 3a1). Although the Mn x MCS x OtoC interaction was not significant (p = 0.269) (Figure 3A), the pattern described by the Mn x OtoC two-way interaction seems to be driven by the fact that MCS diminished the differences between the Mn and control group for distraction trials (Figure 3A); this pattern is not seen in the unsupplemented groups. In light of this, we conducted follow-up analyses where the three trial conditions with olfactory distractors (i.e., OtoC of -1, -2, -3 sec) were collapsed into a single “distraction” trial condition, thus creating only two distraction conditions (distractor and no-distractor trials; Figure 3B). In this follow-up analysis, there was a trending interaction of Mn x MCS x distractor [F(1, 52.16) = 3.29, p = 0.0754]. This trend reflects that, for non-distraction trials, the Mn animals perform more poorly than controls under both diets, but for distraction trials the Mn deficit emerges only for unsupplemented animals (Figure 3B); for the MCS condition, the Mn and controls perform similarly for trials with distractors.

**Figure 3:**
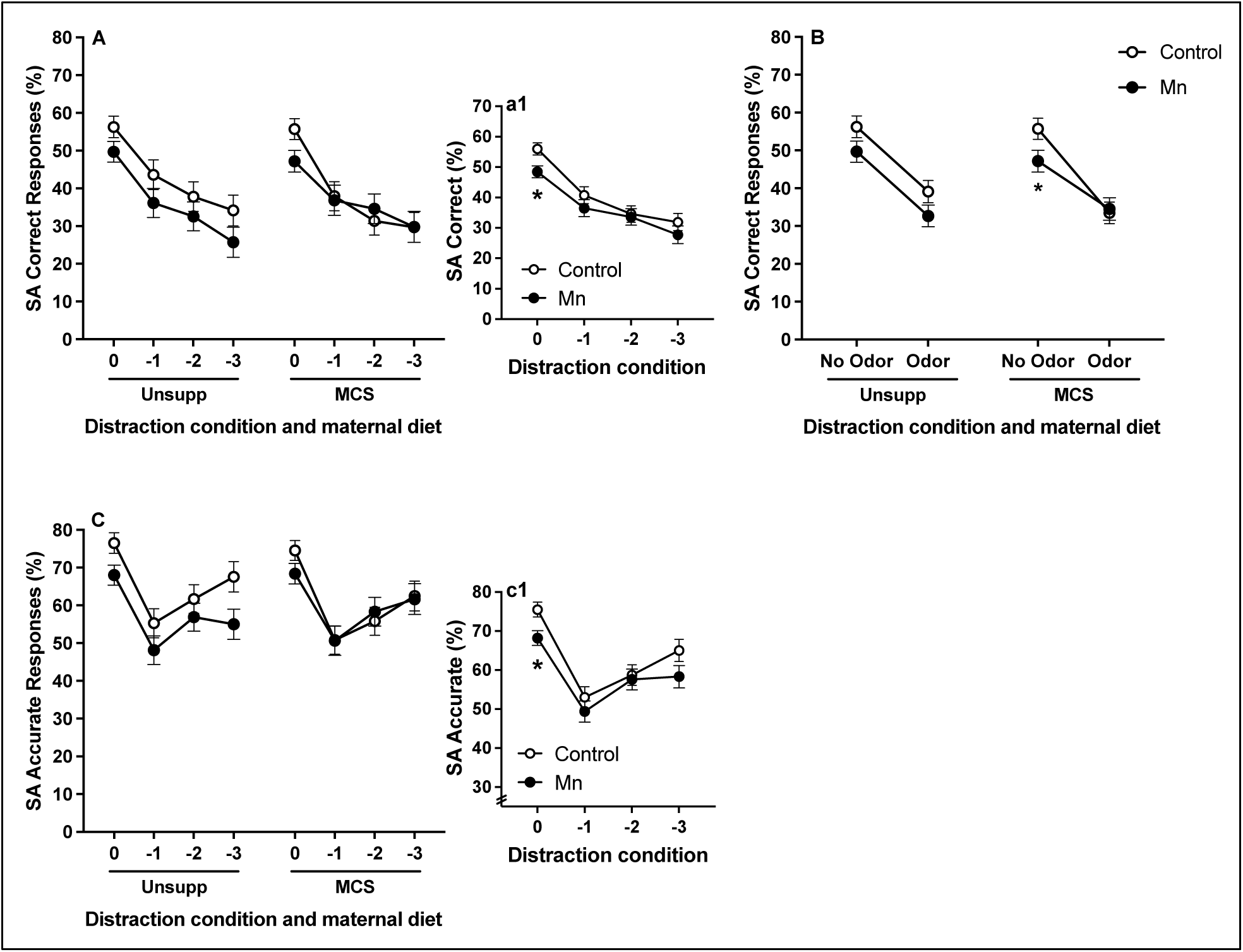
Effects of Mn and MCS on performance in the selective attention task. Panel A depicts %Correct for the analysis in which all four odor distraction conditions were included separately, whereas Panel B depicts %Correct responses for the analysis in which the four odor distraction conditions were collapsed into distractor (odor) vs no distractor (no odor) conditions. Panel C depicts %Accurate responses for all four distraction conditions. Sub-panels a1 and c1 depict significant two-way interactions of Mn x odor distractor condition. Mn causes a deficit in attention, but not selective attention specifically, and MCS reduces the difference between Mn and control animals under distraction conditions. * indicates p ≤ 0.05 versus controls. Data are lsmeans ± SEM (n=14-16/group).

For %Accurate responses, there was a significant Mn x OtoC interaction [F(3, 235.7) = 3.14, p = 0.026], where the Mn deficit was significant for trials without a distractor and trending for trials with the -3 sec distractor condition (p’s = 0.0084 and 0.098; Figure 3c1); the groups were not different for trials with a -1 or -2 distractor. The Mn x MCS x OtoC three-way interaction was not significant [F(3, 235.8) = 0.8, p = 0.495]; however, there is a similar pattern of group differences as noted above for %Correct, where MCS reduces the difference in %Accuracy between control and Mn animals under the distractor conditions (Figure 3C). However, neither the full (Figure 3C) nor distractor-simplified (not shown) %Accurate models revealed a significant Mn x MCS x odor interaction (p’s > 0.31). Collectively, these findings indicate that MCS lessens the difference between control and Mn animals for trials with olfactory distractors.

### 3.4 Behavioral reactivity

#### Mn alters reactivity to errors and reward omission, effects which are partially normalized by MCS

##### In the second focused attention task, Mn animals born to dams on the unsupplemented diet do not exhibit behavioral reactivity to an error, and MCS improves performance in control animals following an error

In the second focused attention task, control animals that received MCS had significantly more correct responses than their unsupplemented counterparts at the 0 sec pre-cue delay (reported above in Figure 2C). We hypothesized that this might be due to MCS reducing the disruptive effect of committing an error on the prior trial, as trials with a 0 sec pre-cue delay are closest temporally to the prior trial (i.e., if the outcome of the prior trial was disruptive to the animal, it would be most apparent for trials for which the cue was presented immediately after trial onset). Therefore, we assessed %Correct responses on trials with a 0 sec pre-cue delay, as a function of the outcome of the prior trial (correct or error).

The analysis of %Correct for trials with a 0 sec pre-cue delay revealed a significant Mn x MCS x previous trial outcome interaction [F(1, 273.2) = 6.13, p = 0.0139]. This interaction reflects that the reaction to committing an error varied as a function of both Mn exposure and maternal diet. Specifically, under the unsupplemented diet, only the controls exhibited fewer correct responses on trials following an error (relative to after a correct response); the Mn animals showed no decline in performance due to making an error on the prior trial (p < 0.0001 for control and p = 0.79 for Mn animals comparing error vs. correct; Figure 4). In contrast, under the MCS condition, both control and Mn groups showed a trending reduction in %Correct for trials following an error (relative to after a correct response; p = 0.074 for control and 0.17 for Mn), and the magnitude of the disruptive effect of making an error was similar in the two groups (Figure 4). Moreover, in the control animals, MCS (vs. the unsupplemented condition) significantly increased correct responses after an error, but not following a correct response (p’s=0.05 and 0.50, respectively), while in the Mn animals, MCS did not improve performance following either a correct response (p = 0.34) or error (p = 0.73), compared to unsupplemented animals (Figure 4).

**Figure 4:**
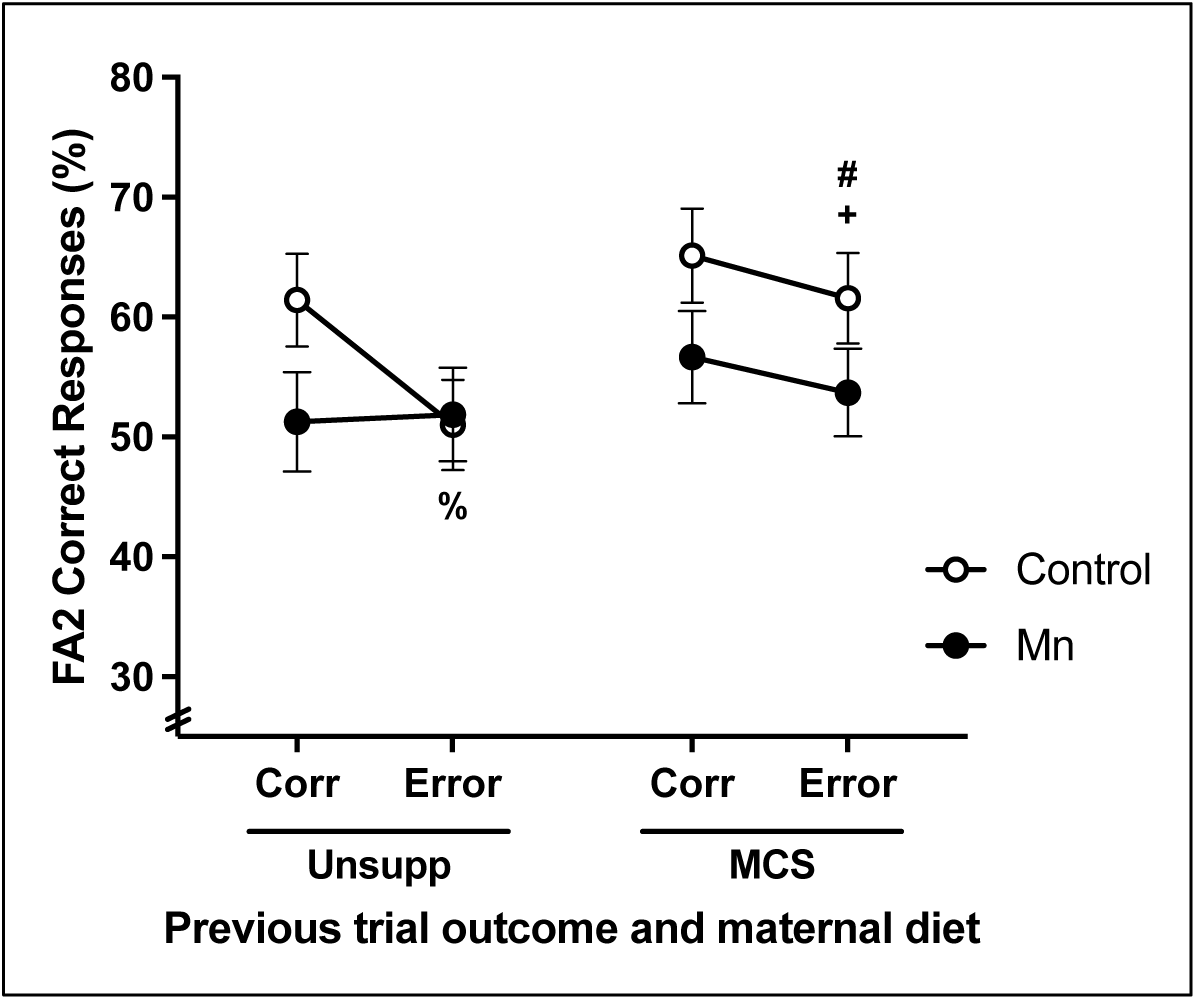
Effects of Mn and MCS on performance in the current trial as a function of the prior trial outcome (i.e., correct or incorrect) in the second focused attention task for trials with a 0 sec pre-cue delay. For animals born to dams on the unsupplemented diet (left side), Mn blunts reactivity to a prior error. Maternal choline supplementation (right) normalizes error reactivity of the Mn animals. Prior trial outcomes are Corr (correct response) or Error (incorrect, premature, or omission error). % and + indicate p ≤ 0.05 and p ≤ 0.10 versus following a correct response, respectively; # indicates p ≤ 0.05 versus unsupplemented animals. Data are lsmeans ± SEM (n=14-16/group).

In contrast to performance on trials with a 0 sec pre-cue delay, there was no Mn x MCS x previous trial outcome interaction for performance on trials with a 3 sec pre-cue delay [F(1, 273.8) = 1.72, p = 0.19] (not shown). The absence of this three-way interaction for trials with a longer pre-cue delay supports the inference that the group differences seen at the 0 sec pre-cue delay (described above) reflects reactivity to the outcome of the prior trial.

### Developmental Mn exposure and MCS both alter reactivity to committing an error and reward omission in the reward omission task

In the reward omission task, we introduced a novel trial condition in which the food pellet reward is unexpectedly omitted on a subset (30%) of correct response trials and assessed how this affects performance on the subsequent trial. Both errors and reward omission have been shown to be disruptive to performance in our prior studies with similar operant tasks^33,67^, and both are often altered in children with ADHD compared to typically developed children^3,4,68–70^. Specifically, we assessed performance on each trial as a function of whether the preceding trial response was: (1) correct with a food pellet reward, (2) an error (incorrect, no food pellet reward + 5 sec time out), or (3) correct but with the food pellet reward omitted (RO).

#### In the unsupplemented animals, Mn lessens reactivity to an error but increases reactivity to the omission of an expected reward

For %Correct responses, there was a significant three-way interaction of Mn x MCS x previous trial outcome [F(2, 410.9) = 4.13, p = 0.0168] (Figure 5A). This reflects that for the unsupplemented controls, committing an error on the prior trial caused a reduction in %Correct on the subsequent trial (versus a correct response on the prior trial; p < 0.0001), whereas the unexpected omission of the food reward on the prior (correct) trial had no effect on performance (p = 0.17). A markedly different pattern was seen in the unsupplemented Mn animals; for these animals, committing an error on the prior trial did not affect performance compared to trials following a correct response (p = 0.152) (Figure 5A). In contrast, performance of the unsupplemented Mn animals was significantly decreased on trials following the omission of the expected food reward relative to after a correct response (p = 0.0032). These findings demonstrate that the unsupplemented animals exposed to Mn early in life respond differently than unexposed controls (no Mn) to both committing an error and not receiving an expected reward.

**Figure 5:**
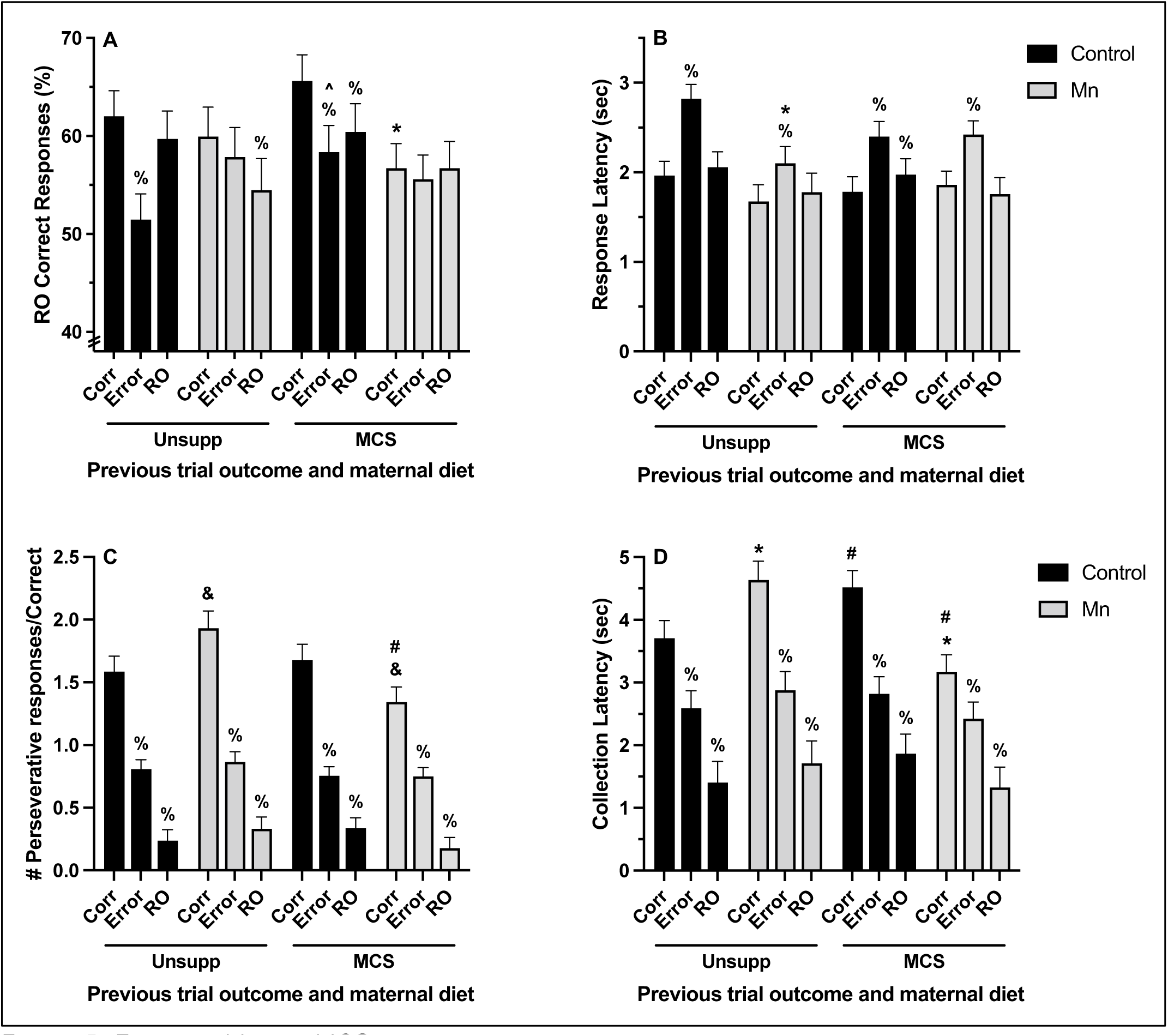
Effects of Mn and MCS on performance in the current trial as a function of the prior trial outcome in the reward omission task. Panels A, B, C and D depict, respectively, %Correct, response latency, perseverative responses, and food pellet collection latency. For the unsupplemented animals, Mn blunts error reactivity, but heightens reactivity to an omitted reward. Maternal choline supplementation (MCS) partially normalizes reactivity to both errors and reward omissions of the Mn animals. Prior trial outcomes are Corr (correct response), Error (incorrect or omission error), or RO (correct with reward omitted). % indicates p ≤ 0.05 versus following a correct response; * and & indicate p ≤ 0.05 and p ≤ 0.10 versus controls, respectively; # and ^ indicates p ≤ 0.05 and p ≤ 0.10 versus unsupplemented animals, respectively. Data are lsmeans ± SEM (n=14-16/group).

#### MCS normalizes behavioral reactivity to an omitted reward in the Mn animals

Notably, a different pattern of performance was seen for the MCS animals. Specifically, for the Mn-exposed animals, MCS normalized performance on trials following a reward omission, such that performance was the same as for trials following a correct response – a pattern matching that of the unsupplemented controls (no Mn) (Figure 5A). However, MCS did not alter how the Mn animals perform following an error (vs a correct response), compared to their unsupplemented Mn-exposed counterparts (Figure 5A). Finally, under the MCS diet, performance on trials following a correct response was significantly worse in the Mn animals than controls (p = 0.0189), a deficit that was not present in the unsupplemented groups; this appears to be because the controls benefited from MCS with improved performance on trials following a correct response, but the Mn animals did not (Figure 5A).

#### For unexposed controls, MCS lessens the disruptive effect of committing an error

For the unexposed controls, there was a trend for MCS to lessen the disruptive effect of committing an error on the prior trial; i.e., to improve performance on trials that followed an error (relative to post-error trials for the unsupplemented controls) (p = 0.076, Figure 5A). However, for the MCS controls, %Correct responses on trials following an error was still lower than on trials that followed a correct response (p < 0.0001). Furthermore, for the MCS controls, performance on trials following a reward omission was lower than that on trials following a correct response (p = 0.0015), an effect that was not seen in the unsupplemented control animals. However, it is notable that performance on trials following a reward omission did not differ between the unsupplemented and supplemented controls; thus, the relative drop in performance by the MCS controls on trials following a reward omission (versus performance on trials after a correct response) was due to an *increase* in performance on trials following a correct response, not a decline in performance following reward omission.

#### Mn animals exhibit less post-error slowing than controls, and MCS normalizes this effect

For response latency, there was a three-way interaction of Mn x MCS x previous trial outcome [F(2, 121.4) = 5.11, p = 0.0074] (Figure 5B). This reflects that for animals born to dams on the unsupplemented diet, both control and Mn groups exhibit post-error slowing (i.e., increased response latency following an error vs. a correct response; p < 0.0001); however, for the unsupplemented groups, the post-error slowing was significantly less pronounced in Mn animals than controls (p = 0.0046), and the magnitude of the difference in response latency following a correct response compared to an error is smaller in the Mn versus control animals (Figure 5B). Maternal choline supplementation normalizes post-error slowing in the Mn animals, such that the response latency following an error is the same as in controls, and the magnitude of the slowing (i.e., increase in response latency following an error versus a correct response) is comparable in the Mn and control animals. Finally, trials following a reward omission have a comparable response latency to trials following a correct response across treatment groups, except in control animals that received MCS; these animals exhibit a slight but statistically significant increase in response latency following a reward omission compared to a correct response (p = 0.044) (Figure 5B).

#### Mn increases perseverative responding following a correct response in the unsupplemented animals, but this abnormality is not seen in the Mn animals born to supplemented dams

The analysis of perseverative responses per correct response revealed a three-way interaction of Mn x MCS x previous trial outcome [F(2, 111.2) = 3.54, p = 0.0322] (Figure 5C). All animals make more perseverative responses on trials following a correct response than on trials following an error or reward omission (p’s < 0.0001), and they make the fewest perseverative responses on trials following a reward omission (compared to trials following either a correct response or error; p’s < 0.0001). For groups in the unsupplemented diet condition, the Mn animals tend to make more perseverative responses following a correct trial than controls (p = 0.0663). However, for the MCS groups, Mn animals trended to have fewer perseverative responses than controls following a correct response (p = 0.053), and there is a significant decrease in perseverative responses following a correct response in the MCS+Mn animals compared to their unsupplemented counterparts (p = 0.0019) (Figure 5C).

#### Mn increases food pellet collection latency; MCS reduces pellet collection latency in Mn animals and increases collection latency in controls, mirroring the effect seen for perseverative responses

For food pellet collection latency (i.e., the time to collect the food pellet reward following a correct response), there was a Mn x MCS x previous trial outcome interaction [F(2, 410.3) = 6.69, p = 0.0014] (Figure 5D). The pattern of effects seen for pellet collection latency was similar to that seen for perseverative responses, as expected, because committing more perseverative responses delays collecting the pellet reward. In general, pellet collection latency is longest on trials following a correct response, and shortest for trials following a correct response with an omitted reward. Unsupplemented Mn animals have a significantly longer collection latency following a correct response compared to controls (p = 0.0265), while under MCS, Mn animals have a significant decrease in collection latency compared to unsupplemented Mn animals following a correct response (p = 0.0005). In addition, Mn animals that received MCS have significantly shorter collection latency than controls following a correct response (p = 0.0006) (Figure 5D). This effect is enhanced by the fact that MCS also increases collection latency in control animals following a correct response, compared to unsupplemented animals (p = 0.039).

### 3.5 Sensorimotor function

#### Early life Mn exposure causes lasting impairment in sensorimotor function that is not altered by MCS

To determine whether MCS can mitigate the lasting sensorimotor impairments caused by developmental Mn exposure, animals were assessed using the Montoya staircase test of skilled forelimb performance. Mn exposure caused lasting impairment in sensorimotor function, as demonstrated by a significant Mn x step level interaction for the number of pellets taken [F(5, 229.3) = 10.66, p < 0.0001], number of pellets eaten [F(5, 232.6) = 4.57, p = 0.0005], %Success [F(5, 231.4) = 3.26, p = 0.0073], and the number of pellets misplaced [F(5, 356.7) = 5.76, p < 0.0001]. Specifically, there was a significant Mn deficit on steps 4 and 5 of the staircase, where the unsupplemented Mn animals ate significantly fewer pellets and misplaced more pellets than their unsupplemented control counterparts (pellets eaten, p’s = 0.0017 and 0.0002 for steps 4 and 5 respectively; pellets misplaced, p’s = 0.0005 and 0.002) (Figure 6a1 and 6b1, respectively). MCS did not affect this Mn-induced impairment in reaching and grasping, based on findings that neither the main effect of MCS (p’s >0.16), nor any interactions involving MCS and Mn were significant for any of the outcomes (e.g., Mn x MCS, P’s >0.32; Mn x MCS x step level, p’s >0.72); see figure panels 6A and 6B for the Mn x MCS x step plots for pellets eaten and misplaced).

**Figure 6:**
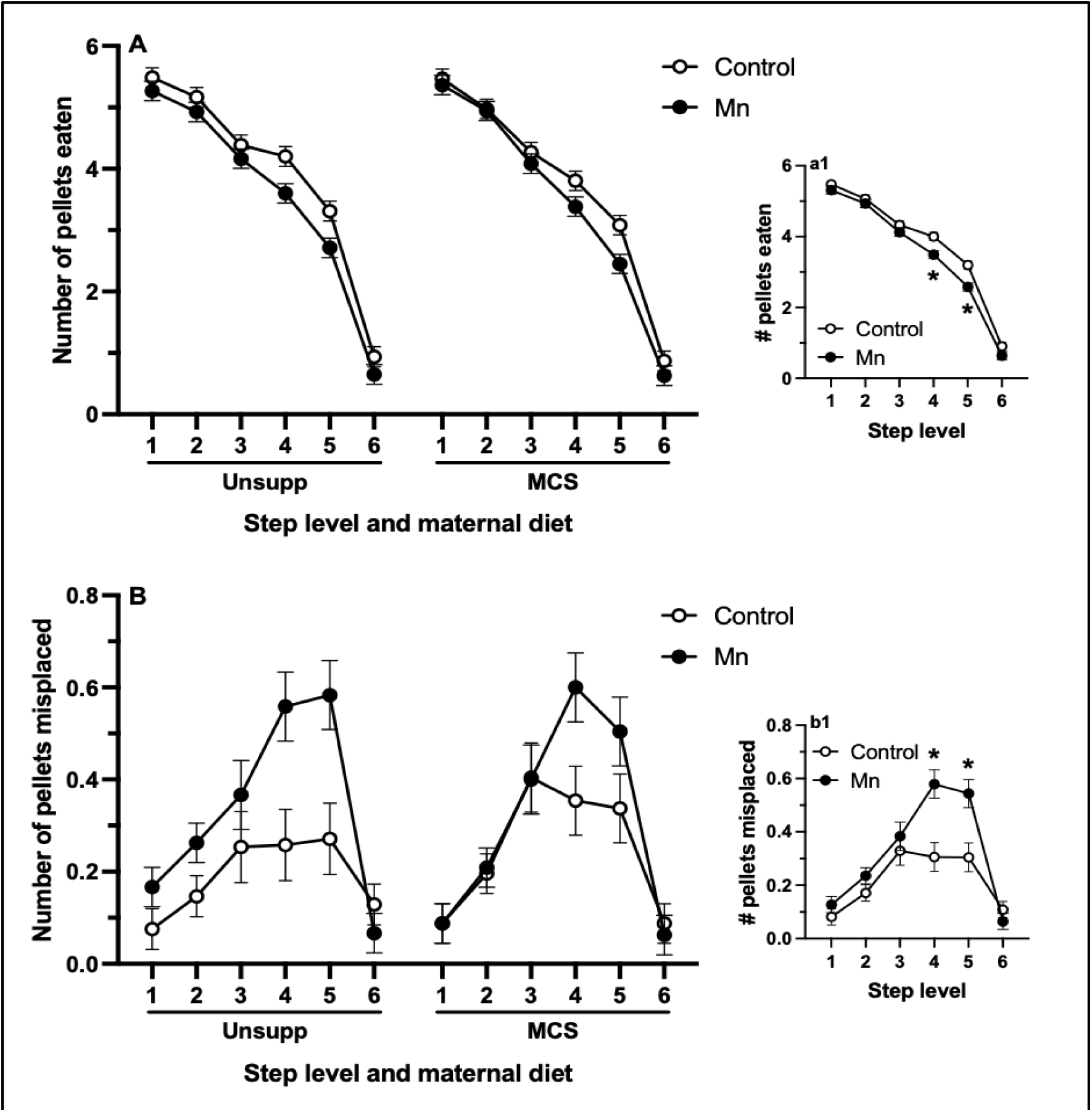
Effects of Mn exposure and MCS on sensorimotor function on the Montoya staircase task. Panel A depicts the number of pellets eaten, whereas Panel B depicts number of pellets misplaced, both as a function of step level. The insert panels show performance collapsed across the two diet conditions. Mn exposure causes a lasting deficit in fine motor function that is not rescued by MCS. * indicates p ≤ 0.05 versus controls. Data are lsmeans ± SEM (n=14-16/group).

### 3.6 Choline and choline metabolite levels

#### MCS measurably altered choline and choline metabolite levels in dams and offspring

To confirm that oral MCS over gestation and lactation altered levels of choline and/or its metabolites in dams and their offspring as expected, and whether oral Mn exposure altered choline/metabolite status in the offspring of MCS-treated dams, we measured levels of choline and selected metabolites in dam and offspring tissue (dam plasma and liver upon completion of MCS at PND 21, offspring liver and brain at PND 24). As expected, MCS produces measurable changes in choline and some of its metabolites in dam plasma and liver, and PND 24 pup liver and brain (see Supplemental Table S2 for choline and metabolite measurements in dams, and Supplemental Table S3 for measurements in pups).

### 3.7 Blood and brain Mn levels

#### Oral Mn dosing increased blood and brain Mn levels in weanlings, and returned to control levels in adulthood

Oral Mn exposure over PND 1-21 increased Mn levels in both blood and brain tissue in PND 24 weanlings. Specifically, Mn exposure significantly increases blood Mn levels ∼eight-fold, from ∼30 to 250 ng/mL in control vs. Mn exposed groups [F(1, 45) = 398.4, p < 0.0001] (Supplemental Figure S4A). There was also a significant interaction of Mn x MCS on blood Mn levels [F(1, 42) = 8.37, p = 0.0059]; however, Tukey’s contrasts revealed no significant difference between blood Mn levels in unsupplemented versus MCS animals exposed to Mn.

Mn exposure also causes an approximately three-fold increase in brain Mn levels in PND 24 offspring, from approximately 3.8 to 11.8 µg/g in control versus Mn-dosed animals, as demonstrated by a significant main effect of Mn [F(1, 36) = 129.7, p < 0.0001] (Supplemental Figure S4B). There is also a trending main effect of MCS on brain Mn levels in offspring [F(1, 36) = 3.96, p = 0.0543], as well as a trending Mn x MCS interaction [F(1, 36) = 3.13, p = 0.085]. Tukey’s pairwise comparisons revealed that MCS animals treated with Mn had significantly higher brain Mn levels than their unsupplemented counterparts.

Finally, while blood and brain Mn levels in the PND 200 behaviorally-tested animals return to baseline levels, there remained a statistically significant but biologically unremarkable effect of MCS [F(1, 27) = 7.87, p = 0.0092] to reduce blood Mn levels compared to the unsupplemented group, and a similar trending effect of MCS to reduce brain Mn levels in the Mn-exposed group [F(1, 20) = 3.19, p = 0.089] (Supplemental Figure S4D). However, Tukey’s pairwise comparisons revealed no significant difference between any groups.

## 4. Discussion

ADHD, which is characterized by impairments in attention, impulse control, and emotion regulation, is the most prevalent neurobehavioral disorder in children, yet the causes of ADHD remain poorly understood^1^. Studies in children and adolescents have revealed associations between developmental Mn exposure and inattention, impulsivity/hyperactivity, emotion dysregulation, and sensorimotor dysfunction^7,8,17,9–16^. However, there are currently no recognized interventions, other than exposure prevention, to mitigate the neurotoxic effects of elevated developmental Mn exposure. Here, using a rodent model of environmental Mn exposure, we demonstrate that developmental Mn exposure causes lasting dysfunction in attention, behavioral reactivity, learning, and sensorimotor function, and that increasing maternal choline intake during pregnancy was effective in protecting against some of these Mn-induced impairments, particularly in the domains of attentional function and behavioral reactivity. MCS also was beneficial for the unexposed animals in the domains of learning, attention, and behavioral reactivity. Collectively, these findings further establish developmental Mn exposure as a significant risk factor for environmentally-induced ADHD and related symptoms. They also demonstrate that MCS exerts lifelong cognitive benefits to offspring, including providing some protection against the neurotoxic impacts of elevated Mn exposure. Given recent evidence that 90% of pregnant women consume less choline than the recommended Adequate Intake (AI), these data provide additional support for efforts to increase choline intake during pregnancy and lactation, particularly for women at risk of environmental exposure to Mn. These findings and inferences are discussed in detail below.

### 4.1 Developmental Mn exposure causes lasting dysfunction in attention, learning, behavioral reactivity, and sensorimotor function, recapitulating the pattern of symptoms reported in children with ADHD

Our findings demonstrate that developmental Mn exposure causes lasting dysfunction in attention, impulse control, sensorimotor function, and behavioral reactivity (i.e., emotion regulation) – a constellation of impairments commonly seen in children diagnosed with ADHD^2–4^. For example, across the three attention tasks, the Mn-exposed animals are particularly impaired in the later session blocks of testing (FA 1 task), and/or under trial conditions with the longest pre-cue delays (FA 2 task). The former finding reflects that the Mn animals improve in attentional accuracy more slowly over testing than controls, while the latter likely reflects deficits in sustained/focused attention at the most challenging (longer) pre-cue delays. Similarly, the Mn exposed animals appear to be more impulsive than controls, based on higher rates of premature responses across all three attention tasks (Figure 1). We should note, however, that the pattern of %Premature responses across session blocks within each attention task for the offspring of unsupplemented dams suggests that the higher rates of premature responses in the Mn animals in some session blocks may be a result of Mn animals learning to withhold premature responses more slowly than controls, rather than inherently greater impulsivity per se. Finally, developmental Mn exposure also causes lasting impairments in sensorimotor function in the Montoya staircase test – impairment that is most pronounced at the steps that are the most challenging to reach, based on significant reductions in the number of pellets taken, pellets eaten, %Success, and increased number of pellets misplaced (Figure 6). Collectively, these findings are fully consistent with our multiple prior studies^18–24^, and they further implicate elevated developmental Mn exposure as a significant risk factor for environmentally-based ADHD and related symptoms^7,8,17,9–16^.

Notably, we also report new findings showing that developmental Mn exposure leads to lasting dysregulation in behavioral reactivity, an essential functional domain comparable to emotion regulation in children^3,4^. Here, we evaluated behavioral reactivity by assessing how attentional performance was affected by either (a) committing an error on the prior trial, or (b) not receiving an expected food pellet reward. As expected from prior studies ^33,67,71–74^, we found that control animals perform significantly more poorly on trials following an error (relative to after a correct response), and that they slow their response time on trials following an error (Figure 4, 5A, B), a phenomenon called “post-error slowing” ^33,67,71–77^. This pattern of changes likely reflects the fact that processing an error diverts attentional resources, leading to both impaired performance and longer response latencies on post-error trials^78^. In contrast, for the Mn animals, committing an error on the prior trial did not affect their %Correct responses, and had only a small effect on their correct response latency (Figure 4, 5A, B). These findings indicate that the Mn-exposed animals may be less attentive to performance errors (i.e., negative feedback), and have diminished capacity to learn from trial history and adjust their behavioral strategy accordingly, suggesting impairment in performance monitoring and reactive control^75^. The pattern seen in the Mn exposed animals on trials following an error – less performance disruption and less post-error slowing than seen in controls – mirrors findings showing that ADHD children are less responsive to committing errors than typically developing children^75^, and exhibit diminished post-error slowing^79,80^. Consistent with these findings, neuroimaging results have revealed that children with ADHD have significantly weaker error-related activation in the salience network than typically developing children^81^, suggesting that ADHD children may be less sensitive to detecting the saliency of their errors.

Interestingly, in contrast to their reduced sensitivity to committing an error, the unsupplemented Mn exposed animals exhibit a heightened reaction to the omission of an expected food reward on the prior trial, relative to that seen in controls (Figure 5A, B). The unsupplemented Mn animals exhibit a significant drop in %Correct performance on trials following a reward omission, compared to their performance after a correct response (Figure 5A, B), a pattern not seen in controls. This finding suggests that the Mn animals are more sensitive to reward (and its omission) than controls, a pattern which is again reminiscent of findings with ADHD children. Specifically, studies show that ADHD children often prefer smaller, more immediate rewards over larger, long-term rewards, are more sensitive than typically-developing children to the positive effects of a reward^68,82^; and are more affected by the loss of an anticipated reward^70^. This pattern thus shows clear parallels with our current findings that the Mn rats exhibit greater drops in performance on trials following a reward omission than do controls.

Consistent with these findings, the Mn-exposed animals commit a higher incidence of perseverative responses on trials following a correct response than do controls. Specifically, while animals across all treatment groups commit more perseverative responses on trials following a correct rewarded response trial than on trials following an error or omitted reward trial, this increase is more pronounced for the Mn animals than controls (Figure 5C), a pattern also seen for food pellet reward collection latency (Figure 5D). These findings provide additional support for the inference that the Mn exposed animals are more sensitive to reward/reinforcement than controls, a pattern also seen in ADHD children (noted above). The findings also support that the Mn animals are more compulsive than controls, a suggestion that is again consistent with the suite of symptoms reported in ∼10 – 25% of ADHD children that are co-morbid for both ADHD and Obsessive Compulsive Disorder (OCD)^83–87^.

Collectively, our findings demonstrate a constellation of behavioral/cognitive impairments in developmentally Mn-exposed animals that are also seen in Mn exposed children and those with ADHD more generally; namely, impairments in attention, impulse control, and sensorimotor function, as well as a reduced response to committing an error and a heightened sensitivity to reward/reward omission^3,4,68,69^. These latter *new* findings relating to Mn-induced dysfunction in behavioral reactivity/emotion regulation and error monitoring are highly relevant clinically, since the ability to maintain emotional and cognitive functioning in the face of unexpected events, as well as to learn from trial history and performance and adjust behavioral strategy accordingly, provides benefits in a variety of situations^81,88,89^. These findings are also clinically relevant in light of evidence showing that emotion dysregulation occurs in ∼25-45% of children and ∼30-70% of adults with ADHD, and that emotional problems have a greater negative impact on well-being and quality of life than other ADHD symptoms such as hyperactivity and inattention^3–5^. Thus, our rat model of environmental Mn exposure, used here and in prior studies^18–24^, exhibits the hallmark areas of dysfunction seen in the ADHD phenotype^2^.

### 4.2 MCS offers some protection against the adverse effects of early developmental Mn exposure

Supplementing the maternal diet with additional choline during pregnancy and lactation offers protection against some of the effects of developmental Mn exposure, although the degree of benefit varied by cognitive domain. Two areas where MCS significantly benefited the Mn animals are in the realms of behavioral reactivity/emotion regulation and attention. In the realm of behavioral reactivity, the pattern of findings indicates that MCS offers protection against the effects of Mn exposure on reward sensitivity and reaction to errors. Specifically, animals born to dams on the choline-supplemented diet and then exposed to Mn in the early postnatal period did not exhibit a drop in performance on trials following a reward omission (similar to the unexposed unsupplemented controls), and very different from the Mn animals born to dams on the unsupplemented diet (Figure 5A). In addition, MCS partially normalized the reaction to committing an error for the Mn animals; this was seen for %Correct in the second focused attention task where the drop in performance after an error was similar for the MCS control and Mn animals. In addition, in the reward omission task, MCS normalizes correct response latency on trials following an error for the Mn animals; that is, the Mn+MCS animals exhibit the same degree of post-error slowing to that seen in the control+MCS animals (Figure 5B). Finally, MCS reduced perseverative responding of the Mn animals in the reward omission task on trials after a correct (rewarded) response trial to levels below the control+MCS animals (Figure 5C), and this effect of MCS was accompanied by a reduction in food pellet reward collection latency (Figure 5D). Together, these findings support that MCS normalized compulsivity and reward omission sensitivity in the Mn animals, and partially normalized error reactivity.

Maternal choline supplementation also offered protection against the detrimental effects of Mn exposure on attentional function. This normalization of attentional function is seen in the second focused attention task, where MCS lessens the impairment of the Mn animals at the longer and more challenging pre-cue delays (Figure 2C and D). Similarly, in the selective attention task, MCS normalizes the performance of the Mn animals on distraction trials, such that they no longer differ from controls (Figure 3). In contrast to the protection provided by MCS for the adverse effects of Mn in the realms of behavioral reactivity and attention, no protection was provided with respect to the impairments seen in learning and sensorimotor function. In the first focused attention task, the Mn animals learned to wait for the visual cue more slowly than controls, and MCS did not alter this Mn effect (Figure 2A, B). Further, the grasping and reaching impairments of the Mn animals born to choline-supplemented dams was as pronounced as that seen in the unsupplemented animals (Figure 6).

Collectively, these findings demonstrate that supplementing the maternal diet with additional choline during pregnancy and lactation partially, but not completely, normalizes Mn deficits in the domains of behavioral reactivity/emotion regulation, compulsivity, and attention, but not learning or sensorimotor function. These neuroprotective effects are consistent with other studies that have shown that MCS lessens the adverse cognitive effects of various developmental insults, including models of various neurological disorders (e.g., Down syndrome. Alzheimer’s disease, autism), toxicant exposure (e.g., alcohol), and other environmental influences (e.g., prenatal stress)^28,29,94–99,30–33,90–93^. For example, MCS has been shown to lessen the functional deficits produced by developmental alcohol exposure in both rodent models ^28,35,96,98–100^, and humans^36,101^. And of particular interest to the present study, MCS has been shown to lessen the attentional dysfunction, as well as to normalize behavioral reactivity to committing an error or not receiving an expected reward in a mouse model of Down syndrome^33^. Collectively, these studies provide additional support for increasing choline intake during pregnancy, particularly for women at risk of toxicant exposure.

### 4.4 MCS benefits cognitive function in offspring not exposed to Mn

In addition to the benefits provided to the Mn exposed animals, supplementing the maternal diet with additional choline was also beneficial for the control animals in the realms of behavioral reactivity, attention, and learning. Specifically, for control animals, MCS lessens the disruptive effect of committing an error on the prior trial (Figure 4, 5A; discussed above in section 4.2), suggesting an improved ability to regulate the behavioral disruption/emotional reaction to committing an error. For attentional function, the MCS benefit in control animals is especially evident at the 0 sec pre-cue delay in the second focused attention task (Figure 2C), an effect which may also partly reflect this reduction in error reactivity because trials with a 0 sec pre-cue delay are closest temporally to the prior trial, some of which were error trials. In addition, trending effects for improved attentional performance were seen for the rate of improvement across blocks in the first focused attention task (Figure 2A, B), as well as for performance in the reward omission task on trials following a correct response (Figure 5A).

Altogether, these findings showing cognitive benefits of MCS in control animals are consistent with the MCS benefits reported in prior studies in humans and animal models^38–41,43,46,47,49^. For example, a number of studies by Meck, Williams and colleagues demonstrate that MCS improves attention and memory in rodents^41–44^. Additionally, studies in humans have demonstrated that increased choline intake during pregnancy produces lasting benefits for offspring attention, memory, and executive functioning ^38–40^. Notably, in one controlled feeding study that followed a cohort of children until 7 years of age, random assignment of mothers to a higher choline intake (vs the AI) during the third trimester produced faster processing speed during infancy^39^, as well as superior memory, sustained attention, and executive functioning at 7 years of age^38^.

### 4.4 Putative neurobiological mechanisms

Our rodent model of developmental Mn exposure recapitulates many of the adverse behavioral symptoms reported in ADHD children. While the neurobiological mechanisms underlying ADHD and related symptoms are not well understood, studies suggest that the combination of multiple environmental and biological risk factors lead to hypofunctioning of the catecholaminergic system within the fronto-cortico-striatal loop and other neural circuits underlying executive attention and emotion regulation^102–106^. Similarly, studies from our lab and others have demonstrated that developmental Mn exposure causes hypofunctioning of the catecholaminergic system in the prefrontal cortex and striatum^22–24^, including reducing stimulated dopamine and norepinephrine release^24,25^, and altered levels of dopaminergic and noradrenergic system proteins^22–24^. The catecholaminergic system in the prefrontal cortex/anterior cingulate cortex has been well-established as critical to executive functioning, including attention, emotion regulation, error monitoring, and behavioral inhibition^107–113^, and thus Mn-induced alterations to this system may well underlie the ADHD-like behavioral phenotype of the Mn exposed animals. This inference is further supported by studies from our group showing that oral methylphenidate (a catecholamine reuptake inhibitor) ameliorates the Mn behavioral deficits in attention and sensorimotor function^19,21,114^. Notably, our group has also demonstrated that developmental Mn exposure causes lasting alterations in DNA methylation and expression of genes related to DNA methylation, neuronal development and catecholaminergic neuronal systems in our rat model^115^, and other studies have also shown that Mn exposure can lead to altered DNA methylation^116–119^. Related to this, it is known that choline serves as, among other things, a methyl donor in key biochemical reactions, and supplementing the maternal diet with additional choline has been demonstrated to alter DNA methylation in both humans and animal models^35,120–126^. Additionally, MCS has been shown to normalize alterations in DNA methylation caused by fetal alcohol exposure^34,35^. Therefore, it is plausible that the benefits of MCS to partially normalize Mn-induced behavioral reactivity and attention deficits in the present study may be due in part to normalization of Mn-induced epigenetic alterations, though this suggestion is in need of investigation.

## 4.5 Conclusion

The present study provides new evidence that developmental Mn exposure produces lasting dysregulation of behavioral reactivity/emotion regulation, in addition to its detrimental effects on attention, impulse control, and sensorimotor function. These findings largely recapitulate the constellation of symptoms reported in Mn-exposed children, and separately in ADHD children, thus further supporting concerns over developmental Mn exposure as a significant risk factor for ADHD and related symptoms. In addition, the present study demonstrates that supplementing the maternal diet with additional choline offers some protection to the adverse effects of early developmental Mn exposure, as well as provides lasting cognitive benefits for the unexposed offspring. In light of the markedly increased physiological demand for choline during pregnancy and the low choline intakes of most pregnant women, these results provide additional support for efforts to increase choline intake during pregnancy, via diet and/or adding choline to standard prenatal vitamin regimens.

## Acknowledgments

The authors would like to thank Bahtia Bilow, Charlyn Agbayani, David Woodfin, Eric Hsu, Gladys Chiang, Jonathan Do, Jordan Altshul, Katie Sharifi, Levon Shahnazarian, Naomi Schlipp, Neha Chenna, Ramtin Poustinchi, Roshana Basto, Sarah Hanson, Sasha (Aleksandra) Nikolaeva, Shannon Twardy, Sofia Bayliss Grijalva, Tyler Waite, Vanessa Howland, Victoria Deng, and Wendy Valdez Rodriguez for their invaluable assistance in behavioral testing. We also thank Tom Jursa, Dr. Marie Caudill, and Olga Malysheva for analytical assistance, Dr. Lynn Johnson for assistance with statistical analyses, and Dr. Nicholas Santiago for assistance in the study.

## Author contributions

D.R.S., S.A.B., B.J.S., and S.L.H. conceived and designed research. S.L.H., S.A.B., and D.R.S. performed experiments. S.L.H., S.A.B., B.J.S., and D.R.S. analyzed and interpreted results of experiments. S.L.H. prepared figures. S.L.H., S.A.B., B.J.S., and D.R.S. drafted manuscript. S.L.H., S.A.B., B.J.S. and D.R.S. edited and revised manuscript.

