## Supplemental Tables/Figures for "Maternal choline supplementation lessens the behavioral dysfunction produced by developmental manganese exposure in a rodent model of ADHD"

Figure S1. Study timeline with task parameters, and study design.

Table S2. MCS produces measurable changes in choline and some of its metabolites in dam plasma and liver.

Table S3. MCS and Mn produce measurable changes in choline and some of its metabolites in PND 24 pup liver and brain.

Figure S4. Oral Mn dosing increased blood and brain Mn concentrations in weanling littermates of the behaviorally tested animals (PND 24), and returned to baseline in the behaviorally tested animals at sacrifice (PND 200).

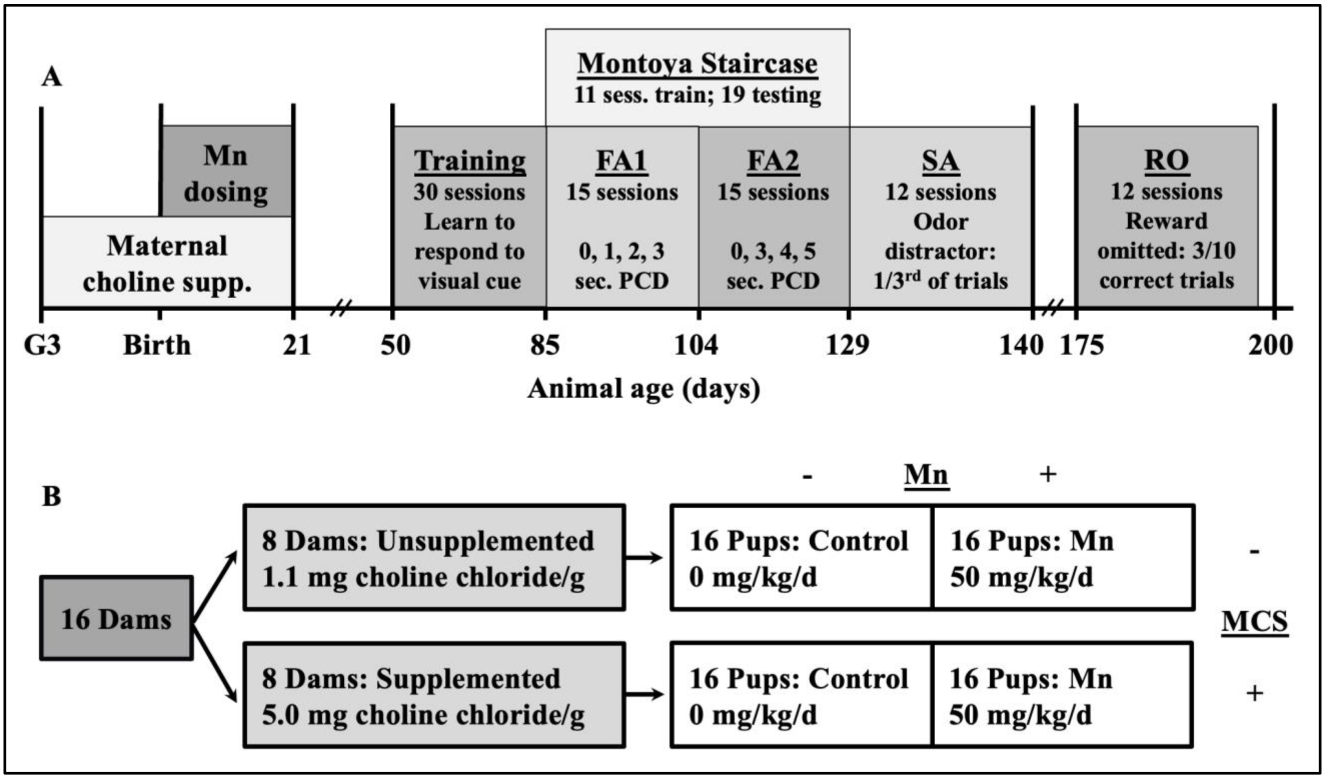


**Figure S1. Study timeline with task parameters, and study design.** (A) Study timeline, including MCS and Mn dosing, as well as tasks with parameters. X-axis is days of age (PND). (B) 2x2 study design, ending with 16 pups per treatment group that were used for behavioral testing.

| Dam Tissue | Metabolite | Unsupp. diet | MCS diet | MCS effect |
|---|---|---|---|---|
| Plasma nmol/mL | Dimethylglycine (DMG) | 1.57 ± 0.18 | 3.36 ± 0.40 * | p = 0.0020 |
| | Betaine | 78.6 ± 4.6 | 174 ± 17 * | p = 0.0002 |
| | Trimethylamine N-oxide (TMAO) | 1.40 ± 0.18 | 34.0 ± 11 * | p = 0.0178 |
| | Choline | 21.3 ± 4.0 | 41.4 ± 8.6 # | p = 0.0638 |
| Liver nmol/g | Choline | 98.2 ± 22 | 371 ± 100 * | p = 0.0185 |
| | Methionine | 125 ± 7.7 | 87.6 ± 11 * | p = 0.0145 |
| | Betaine | 199 ± 18 | 662 ± 130 * | p = 0.0034 |
| | TMAO | 2.54 ± 0.44 | 70.0 ± 28 * | p = 0.0322 |
| | Lysophosphatidylcholine (LPC) | 445 ± 14 | 524 ± 13 * | p = 0.0011 |
| | DMG | 15.6 ± 1.3 | 31.9 ± 6.8 * | p = 0.0333 |
| | Glycerophosphocholine (GPC) | 536 ± 46 | 807 ± 140 # | p = 0.0909 |

**Table S2: MCS produces measurable changes in choline and some of its metabolites in dam plasma and liver.**

MCS increases plasma DMG, betaine, and TMAO, and trends toward increasing choline levels, while MCS increases liver choline, betaine, TMAO, LPC, and decreases liver methionine. MCS also trends toward increasing GPC in liver. * indicates $p \leq 0.05$ versus unsupplemented diet; # indicates $p \leq 0.10$ versus unsupplemented diet. Statistical analyses used standard least squares models. Data are mean ± SEM (n=8/group).

**Metabolites measured include:**

**Dam plasma**: methionine, choline, betaine, dimethylglycine (DMG), Trimethylamine N-oxide (TMAO)

**Dam liver**: methionine, choline, betaine, DMG, TMAO, glycerophosphocholine (GPC), phosphocholine, phosphatidylcholine (PtdCho), sphingomyelin (SM), lysophosphatidylcholine (LPC)

| Pup Tissue | Metabolite | Control Unsupp. diet | Control MCS | Mn Unsupp. diet | Mn MCS | Mn main effect | MCS main effect | MCS*Mn interaction |
|---|---|---|---|---|---|---|---|---|
| Brain nmol/g | GPC | 315 ± 15 | 488 ± 80 | 349 ± 18 | 400 ± 36 | p = 0.513 | *** *p = 0.0159*** | p = 0.158 |
| | LPC | 166 ± 7.7 | 177 ± 7.1 | 156 ± 11 | 185 ± 12 | p = 0.903 | **# p = 0.0570** | p = 0.386 |
| | Sphingomyelin (SM) | 5950 ± 290 | 5330 ± 150 | 5180 ± 280 | 4990 ± 280 | **# p = 0.0557** | p = 0.147 | p = 0.437 |
| Liver nmol/g | Lysophosphatidylcholine | 352 ± 13 | 364 ± 5.1 | 330 ± 5.9 | 361 ± 10 | p = 0.205 | *** *p = 0.0384*** | p = 0.324 |
| | Betaine | 3750 ± 170 | 2610 ± 230 | 3870 ± 480 | 3450 ± 330 | p = 0.176 | *** *p = 0.0361*** | p = 0.307 |
| | SM | 1610 ± 29 | 1450 ± 34 | 1450 ± 20.5 | 1190 ± 33 | *** *p < 0.0001*** | *** *p < 0.0001*** | p = 0.158 |
| | Phosphatidylcholine | 20100 ± 630 | 21200 ± 490 | 19400 ± 390 | 19800 ± 120 | *** *p = 0.0345*** | p = 0.130 | p = 0.384 |

**Table S3: MCS and Mn produce measurable changes in choline and some of its metabolites in PND 24 pup liver and brain.** In offspring, MCS increases GPC in brain, trends toward increasing LPC in brain, increases LPC in liver, and decreases betaine and SM in liver. Oral Mn exposure also alters levels of some metabolites in PND 24 pups, including trending toward decreasing sphingomyelin in brain, and decreasing SM and PtdCho in liver. Data are means ± SE. Statistical analyses used standard least squares models. * indicates a main treatment effect of p ≤ 0.05; # indicates a main treatment effect of p ≤ 0.10.

**Metabolites measured include:**

**Pup liver**: methionine, choline, betaine, GPC, phosphocholine, PtdCho, SM, LPC

**Pup brain**: methionine, choline, betaine, GPC, phosphocholine, PtdCho, SM, LPC, and acetylcholine

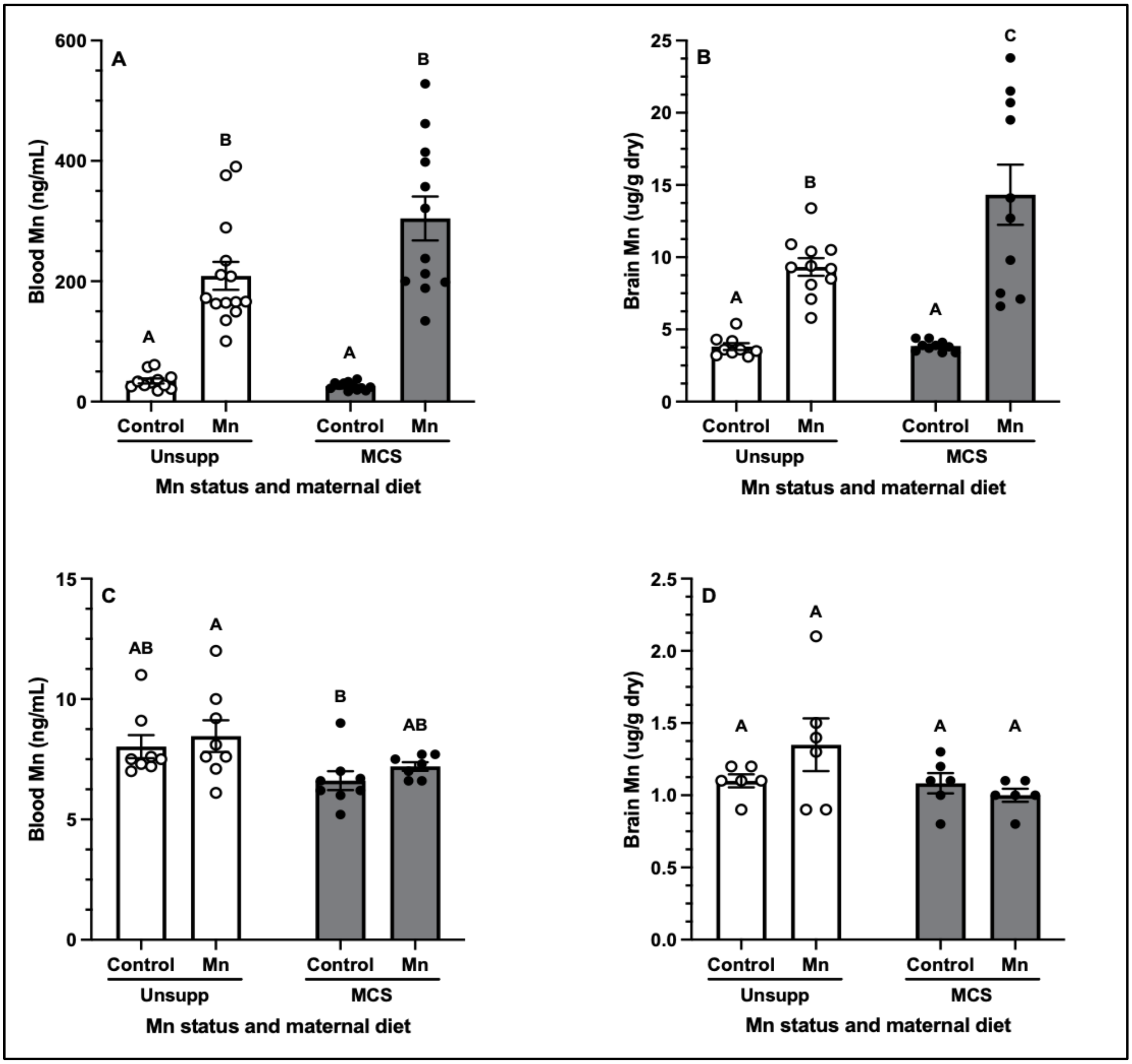


**Figure S4: Oral Mn dosing increased blood and brain Mn concentrations in weanling littermates of the behaviorally tested animals (PND 24), and returned to baseline in the behaviorally tested animals at sacrifice (PND 200).** PND 24 animal blood (n = 11-14) (A) and brain (n = 9-11) (B) measurements show that Mn was increased in Mn-exposed animals. PND 200 animal blood (n = 7-8) (C) and brain (n = 6) (D) measurements show that Mn levels return to baseline. Graphs show arithmetic means with standard error. PND 24 blood and brain statistics were performed on log-transformed data; PND 200 statistics were performed on non-transformed data. Within an age group and tissue, treatment groups with different capital letter superscripts are statistically different from one another ($p < 0.05$), based on Tukey's post hoc test.
